# Mechanisms of permselectivity of connexin hemichannels to small molecules

**DOI:** 10.1101/2025.03.12.642803

**Authors:** Alexandra Lovatt, Jack Butler, Nicholas Dale

## Abstract

Connexins can either that act as hemichannels, to facilitate ion and small molecule movement from the cytosol to the extracellular space or as gap junction channels to provide a pathway for solute exchange between cells. Connexins are ubiquitously expressed throughout the body and are implicated in a wide range of processes. The permselectivity of connexin hemichannels for small neurochemicals remains poorly understood. By coexpressing genetically encoded fluorescent sensors for ATP, glutamate and lactate with a range of connexins, we examined the ability of different hemichannels to permit release of these compounds under physiological conditions and in response to physiological stimuli (small changes in PCO^2^ and transmembrane depolarisation). We found that some connexin hemichannels were relatively non-selective (Cx26, Cx32, Cx43, Cx31.1) allowing passage of ATP, glutamate and lactate. By contrast other connexin hemichannels (Cx36, Cx46 and Cx50) were highly selective. Cx36 and Cx46 hemichannels allowed release of ATP, but not glutamate or lactate. The size of the permeating molecule cannot be the sole determinant of permselectivity. By contrast, Cx50 hemichannels permitted the release of lactate and glutamate but not ATP. We also found that the nature of the opening stimulus could alter the permselectivity of the hemichannel-for some of the relatively non-selective connexins, hemichannel opening via depolarisation was ineffective at allowing release of lactate. By performing a mutational analysis, informed by the differential selectivity of the closely related Cx46 and Cx50 hemichannels, we found that the charge on the N-terminus and N-terminus-TM2 interactions are key contributors to permselectivity for ATP.

## Introduction

There are 21 connexin genes in the human genome (Lucaciu et al., 2023). Connexins form hexamers that, if unopposed, can act as a plasma membrane hemichannel that opens to the extracellular space. However, hemichannels of closely apposed cells can also dock together to form gap junction channels to provide an aqueous passageway between cells. The structure of the hemichannel is highly conserved in all 21 isoforms; each connexin has an N-terminal helix, 4 transmembrane helices, a cytoplasmic loop, 2 extracellular loops and a cytoplasmic C-terminus (Maeda et al., 2009; Myers et al., 2018; Flores et al., 2020; Brotherton et al., 2022; Lee et al., 2023; Qi et al., 2023; Brotherton et al., 2024). 6 subunits then co-assemble to form a hemichannel with a central pore, spanning ∼1.2 nm that is permeable to small molecules up to a molecular weight of about 1000 Da. Major differences in structure between isoforms lie within the cytoplasmic loop and C-terminus, which vary in sequence and length (Mese et al., 2007). The 6 N-terminal helices line the hemichannel pore, to form the narrowest part of the permeation pathway, suggesting that the N-terminus may be an important for determining the permeability of the channel (Oshima et al., 2007, 2008; Maeda et al., 2009; Nielsen et al., 2019; Yue et al., 2021).

Connexin hemichannels have been documented to release small molecules such as ATP under physiological conditions (Weissman et al., 2004; Pearson et al., 2005; Huckstepp et al., 2010b; Chever et al., 2014; van de Wiel et al., 2020). Yet the mechanisms that control hemichannel permeability to different molecules, and whether there is specificity to which molecules may permeate is still unclear. Traditionally, this has been investigated using various fluorescent dyes such as ethidium bromide, and the size and charge of dyes provided some evidence for selectivity to release of larger molecules (Li et al., 1996; Saez et al., 2010; Johnson et al., 2016). Investigation of hemichannel permeability via dye fluxes, while valuable, may differ from how physiological metabolites such as ATP, glutamate or lactate permeate these channels. Traditionally, connexin permeability studies have used the removal of extracellular divalent cations to unblock the hemichannels (Hansen et al., 2014; Nielsen et al., 2019). The mechanism was defined in Cx26 and Cx32 and involves a ring of 12 aspartate residues within the extracellular loop that provide a carboxylate cluster able to bind Ca^2+^ ions with millimolar affinity (Gomez-Hernandez et al., 2003; Bennett et al., 2016; Lopez et al., 2016). While hemichannels are essentially blocked at Ca^2+^ concentrations over 1 mM, there are very few if any physiological conditions in which extracellular Ca^2+^ is lower than 1 mM. Thus, unblocking of hemichannels via Ca^2+^ removal may open a permeation pathway that is not representative of physiological gating of connexin hemichannels. Nevertheless, this method has been used to suggest differential permeability of Cx30 and Cx43 hemichannels to a variety of small molecules (Hansen et al., 2014; Nielsen et al., 2019).

Connexin hemichannels can be opened under physiological conditions by other gating stimuli. As the N-termini of connexins have charged residues and are within the membrane electric field, almost all connexin hemichannels can be opened by sufficient depolarisation, without the need to lower extracellular Ca^2+^ (Pinto et al., 2016). A subset of connexins is directly sensitive to the concentration of gaseous CO^2^ and can be opened by relatively small changes in the partial pressure of CO^2^ (PCO^2^) around the physiological norm. CO^2^-dependent gating was elucidated in Cx26 (Meigh et al., 2013): K125 is carbamylated by CO^2^, which facilitates the formation of a salt bridge with R104 of the adjacent subunit to bias the channel into an open conformation. This mechanism involves movements of the N-terminus, along with related movements of the transmembrane helices (particularly TM2) (Brotherton et al., 2022; Brotherton et al., 2024). CO^2^ gating has subsequently been discovered in a subset of connexins: Cx30, Cx32 (Huckstepp et al., 2010a), Cx43 (Nijjar et al., 2025) and Cx50, since they all have the carbamylation motif that is required for CO^2^-mediated opening. Their CO^2^ sensitivity lies close to the physiological range of PCO^2^; Cx26, Cx43 and Cx50 hemichannels are maximally open at ∼55 mmHg PCO^2^, and Cx32 at ∼70 mmHg. All of these hemichannels are essentially shut at a PCO^2^ of 20 mmHg.

To permit the study of the permeation of connexin hemichannels, for physiological molecules under their normal electrochemical gradients and with physiological gating stimuli (voltage and PCO^2^), we have developed an assay to allow real-time imaging of analyte release at single cell resolution. We utilised the genetically encoded sensors GRAB^ATP^ (Wu et al., 2022), iGluSnFR (Marvin et al., 2013) and eLACCO1.1 (Nasu et al., 2021) to measure the release of ATP, glutamate and lactate respectively via coexpressed connexin hemichannels. We find that connexins can be divided essentially into relatively non-selective and highly selective categories. Connexin hemichannels such as those formed by Cx26, Cx32 and Cx43 fall into the relatively non-selective category, whereas hemichannels composed of Cx36, Cx46 and Cx50 are highly selective. By comparing the highly homologous Cx46 and Cx50, we have shown that key residues in the N-terminus and in the interacting portion of TM2 determine the permeability profile of the hemichannel.

## Results

We first transfected HeLa DH cells with the genetically encoded sensors on their own to ensure that the sensors had no intrinsic responses to CO^2^ or high KCl solutions, or that parental HeLa cells exhibited ATP, glutamate or lactate release in the absence of connexin expression. The median change in normalised fluorescence (ΔF/F^0^) for GRAB^ATP^ was-0.007 (95% CI: 0.0043, - 0.013) and-0.004 (0.0017,-0.0064) for 55 mmHg and 50 mM KCl respectively. GRAB^ATP^ was functional as it gave a median response of 0.2 (0.25, 0.16) to 3 μM ATP (Fig.1). The median change in ΔF/F^0^ for iGluSnFR with 55 mmHg was 0 (0.0008,-0.0026), for 50 mM KCl it was-0.014 (-0.011,-0.030), and for 3 μM glutamate was 0.1 (0.13, 0.05). Finally, the median ΔF/F^0^ for eLACCO1.1 (modified by insertion into iGluSnFR backbone) was 0.005 (0.0073,-0.0046),-0.002 (0.0048,-0.0069) and 0.04 (0.046, 0.032) for 55 mmHg, 50 mM KCl and 3 μM lactate, respectively. The negative recorded values are an artefact of photobleaching. As there were no responses of any of the genetically sensors to a change in PCO^2^ or membrane depolarisation, we conclude that parental HeLa DH cells do not express any channels capable of releasing ATP, glutamate or lactate to these stimuli.

**Figure 1.**
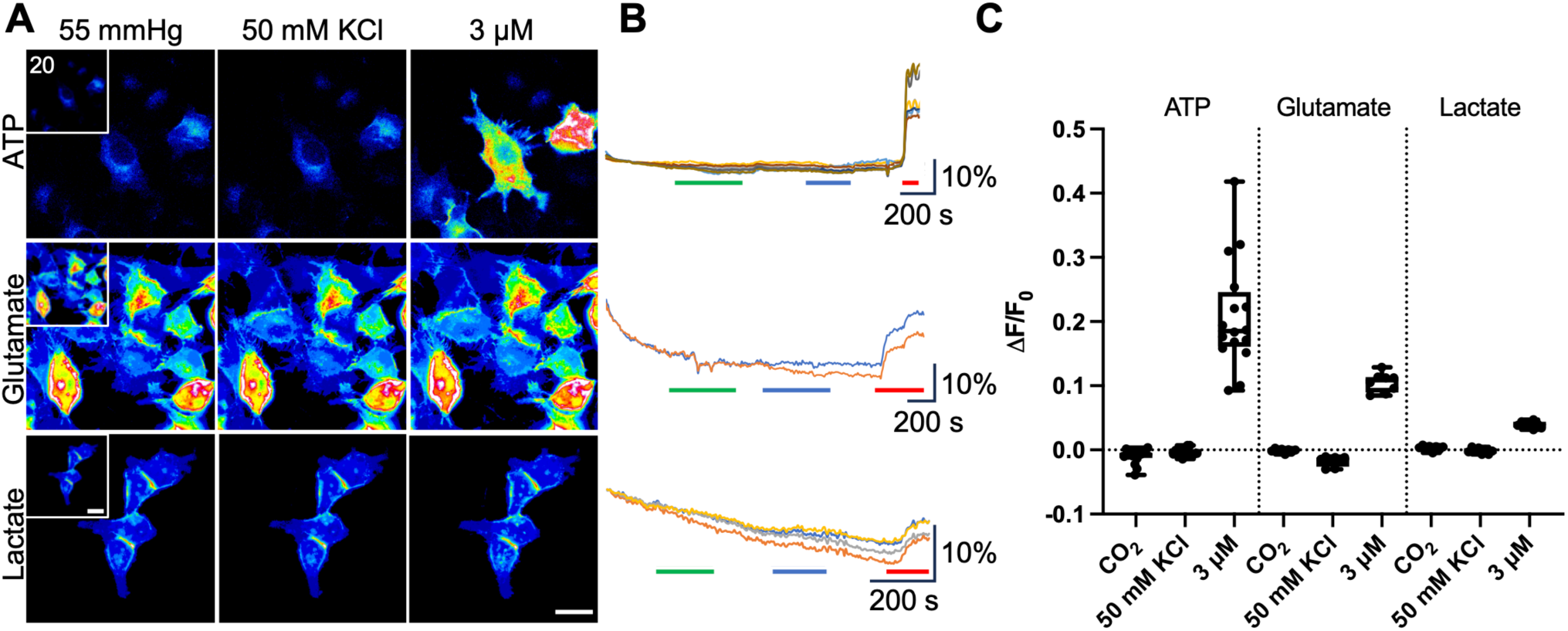
HeLa cells expressing the genetically encoded fluorescent sensors GRAB^ATP^, iGluSnFr and eLACCO1.1 alone do not respond to connexin gating stimuli. A, Representative images of cells at 55 mmHg PCO^2^ (inset is 20 mmHg baseline control), 50 mM KCl, and after application of 3 μM corresponding analyte. Scale bar represents 20 μm. **B,** Representative traces showing sensor responses to 50 mM KCl (green bar), 55 mmHg (blue bar), and 3 μM corresponding analyte (red bar). Vertical scale bars represent the % change in ΔF/F^0^. **C,** Summary data showing median ΔF/F^0^ for ATP (n = 16 cells), glutamate (n = 8) and lactate (n = 8). Box and whisker plots with superimposed data points, showing the median (line), the interquartile range (box) and range (whiskers).

### Cx26, Cx32,Cx43 and Cx31.3 hemichannels are permeable to ATP, glutamate, lactate

We next co-transfected HeLa cells with Cx26, Cx32, Cx43 or Cx31.3 and one of the genetically encoded fluorescent sensors. Throughout all of the assays, we selected cells that co-expressed the connexin and the sensor for measurement and analysis (Supplementary Fig. 1). Cx26 hemichannels are highly permeable to ATP, glutamate and lactate (Fig. 2). The median ATP release to hypercapnic stimuli was 1.5 μM (1.91, 1.32) and with a depolarising stimulus was 2.5

**Figure 2.**
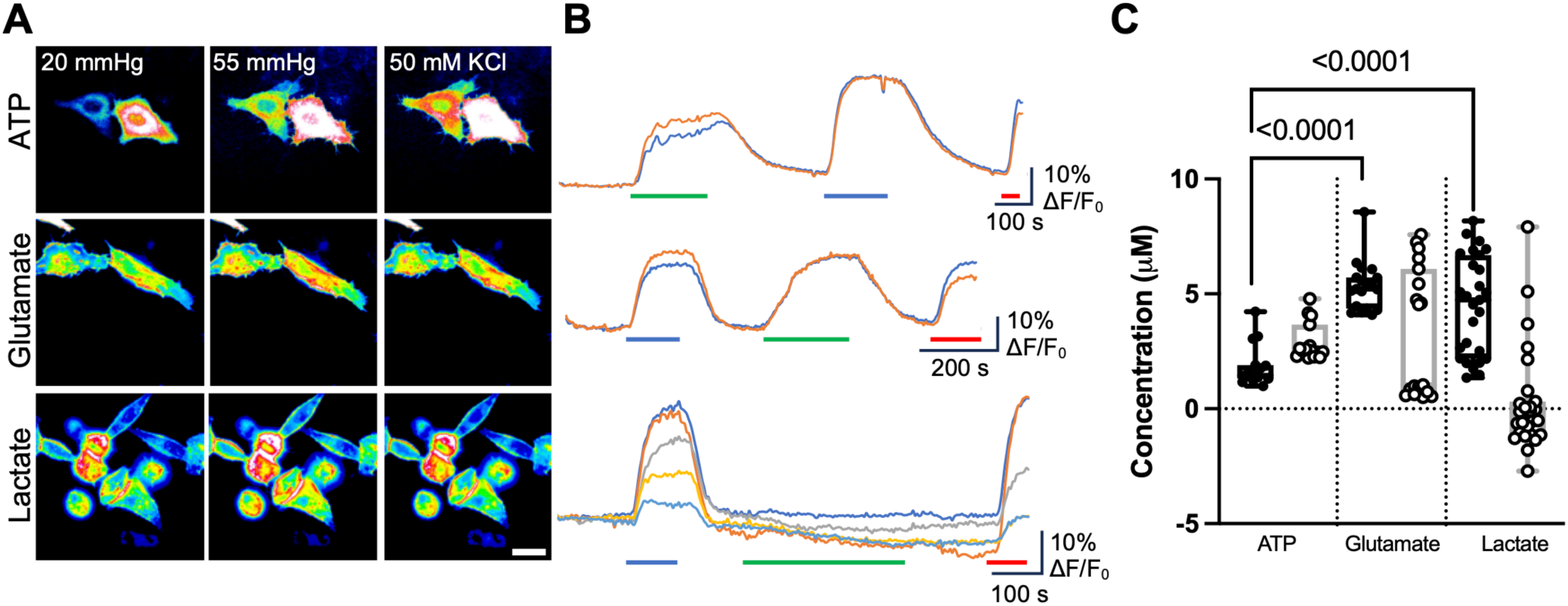
Cx26 hemichannels mediate release of ATP, glutamate and lactate. A, Representative images showing cells from each sensor under each condition. Scale bar represents 20 μm. **B,** Representative traces of normalised fluorescence changes in response to 55 mmHg PCO^2^ (blue bar), 50 mM KCl (green bar) and a 3 μM calibration of the corresponding analyte (red bar). **C,** Summary data showing the median release for ATP (n = 15 cells), glutamate (n = 19) and lactate (n = 27) through Cx26 hemichannels stimulated either by 55 mmHg (filled circles), or 50 mM KCl (open circles). Kruskal Wallis Anova p<0.0001 for 55 mmHg CO^2^ and p<0.0001 for 50 mM K^+^. Mann Whitney U-tests, p < 0.0001 (55 mmHg ATP vs glutamate, 55 mmHg ATP vs lactate). The order of stimulus (CO^2^ or KCl) was regularly reversed between recordings to avoid any potential depletion of ATP release. Data present is from at least 3 independent transfections. Box and whisker plots with superimposed data points, showing the median (line), the interquartile range (box) and range (whiskers).

μM (3.67, 2.30). This was significantly less than for glutamate and lactate: the median glutamate release was 5.2 μM (5.71, 4.36) and 1.1 μM (6.10, 0.74) when opened with hypercapnia and a depolarising stimulus, respectively. Lactate release evoked by hypercapnia was comparable to glutamate: median release of 4.7 μM (6.26, 2.52). However when stimulated by depolarisation no lactate release was evident: median-0.1 μM (0.23,-1.04). For voltage-gated glutamate release it appeared that in some cells the depolarising stimulus was ineffective at evoking release. This suggests that KCl evoked depolarisation (estimated to be about 70 mV from the Nernst equation) is a less reliable gating mechanism than CO^2^ for Cx26.

We determined that Cx32 hemichannels were also permeable to all three tested analytes. Significantly more glutamate and lactate was released compared to ATP regardless of the stimulus (Fig. 3). However the nature of the stimulus did alter the relative amounts of release of the three metabolites. The median release of ATP was 1.3 (1.51, 1.16) and 3.0 (3.12, 2.86) μM when stimulated by hypercapnia and voltage, respectively. Glutamate released via Cx32 hemichannels by hypercapnia and depolarisation was respectively 3.9 (4.64, 3.41) μM and 5.2 (7.44, 4.62) μM. Lactate release via Cx32 hemichannels by hypercapnia and depolarisation was respectively 7.2 (8.32, 5.31) μM and 5.5 (6.84, 4.52) μM. With hypercapnia, the relative release of both glutamate and lactate compared to ATP was considerably more than might be expected from the calculated electrochemical driving force (Tables 1 and 2). This suggests that Cx32 hemichannels may have enhanced permeability for these molecules when opened by hypercapnia.

**Figure 3.**
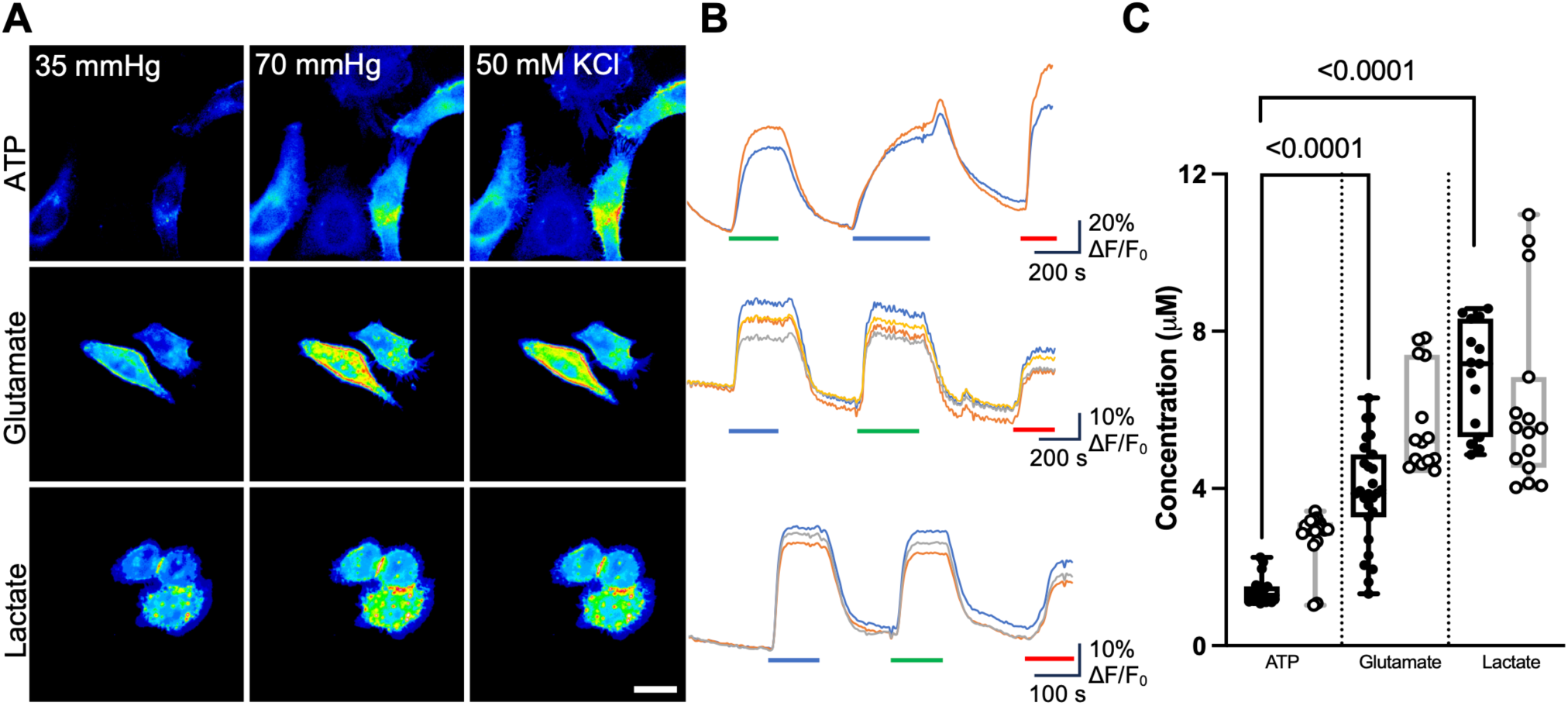
Cx32 hemichannels mediate release of ATP, glutamate and lactate. A, Representative images showing cells from each sensor under each condition. Scale bar represents 20 μm. **B,** Representative traces of normalised fluorescence changes in response to 70 mmHg PCO^2^ (blue bar), 50 mM KCl (green bar) and a 3 μM calibration of the corresponding analyte (red bar). **C,** Summary data showing the median release of: ATP (n = 18), glutamate (n = 27 for CO^2^, n = 14 for KCl), and lactate (n = 15), through Cx32 hemichannels showing changes in release depending on whether the channel was opened by CO^2^ (open circles), or a depolarising stimulus (closed circles). Data presented is from at least 3 independent transfections. Kruskal Wallis Anova p<0.0001 for 55 mmHg CO^2^ and p<0.0001 for 50 mM K^+^. Mann Whitney U-tests, p < 0.0001 (CO^2^ ATP vs glutamate), p < 0.0001 (CO^2^ ATP vs lactate), p < 0.0001 (50 mM KCl ATP vs glutamate), p < 0.0001 (50 mM KCl ATP vs lactate). Box and whisker plots with superimposed data points, showing the median (line), the interquartile range (box) and range (whiskers).

**Table 1.** Calculations of the Nernstian equilibrium potential for ATP, glutamate and lactate in HeLa cells. We assigned the valence of ATP under the assumption that it is chelated to Mg^2+^. Driving force is calculated assuming a resting potential of-60 mV. The intracellular concentrations were taken from studies that investigated HeLa cells: a, (Imamura et al., 2009); b, (Piva and McEvoy-Bowe, 1998); c, (San Martin et al., 2013).

| Analyte | Valence | [Analyte]<sub>i</sub><br>(mM) | [Analyte]<sub>o</sub><br>(mM) | V<sub>eq</sub> (mV) | Driving force at rest |
| --- | --- | --- | --- | --- | --- |
| ATP | -2 | 10<sup>a</sup> | 10<sup>-4</sup> | 145 | 205 |
| Glutamate | -1 | 20<sup>b</sup> | 10<sup>-4</sup> | 307 | 367 |
| Lactate | -1 | 1<sup>c</sup> | 10<sup>-4</sup> | 232 | 292 |

**Table 2.** Relative release of ATP, glutamate and lactate of connexin hemichannels normalised to amount of ATP release. The numbers given are for the hypercapnic stimulus, except for Cx31.3, Cx36 and Cx46 which are insensitive to CO^2^. For those channels sensitive to CO^2^, the numbers in brackets are normalised values for the depolarising stimulus. For Cx50, which is not permeable to ATP, the numbers are absolute concentration in µM. The ratio of the Nernstian driving force for each analyte (normalised to that of ATP) during the hypercapnic stimulus is 1:1.8:1.4, during the depolarising stimulus it is (1:2.1:1.6) assuming that the membrane depolarises to 0 mV.

| Connexin | ATP | Glu | Lac |
| --- | --- | --- | --- |
| Cx26 | 1 | 3.5 (0.44) | 3.1 (0) |
| Cx32 | 1 | 3.0 (1.73) | 5.5 (1.83) |
| Cx43 | 1 | 2.4 (0.4) | 2.0 (0) |
| Cx31.3 | 1 | 0.9 | 1.1 |
| Cx36 | 1 | 0 | 0 |
| Cx46 | 1 | 0 | 0 |
| Cx50 | 0 | 1.5 (0) μM | 5.8 (3.4) μM |

With a hypercapnic stimulus hemichannels composed of Cx43 were permeable to ATP, glutamate and lactate (Fig. 4). The median ATP, glutamate and lactate release were respectively 2.2 (2.34, 1.49) μM, 5.3 (5.71, 4.36) μM and 4.5 (5.84, 2.46) μM (Fig. 4). This amount of release is roughly proportional to the electrochemical driving force on these molecules (Tables 1 and 2) suggesting no selectivity. However during membrane depolarisation evoked by 50 mM KCl, the permeation profile was different. Whereas median ATP release was 2.5 (2.88, 2.22) μM (similar to that evoked by CO^2^), the release of glutamate was reduced, median 1.0 (6.10, 0.74) μM, and lactate was not released at all during this stumulus, median-1.0 (-0.069,-1.49) μM. The gating mechanism of Cx43 thus alters the permeability profile of the hemichannel to small molecules.

**Figure 4.**
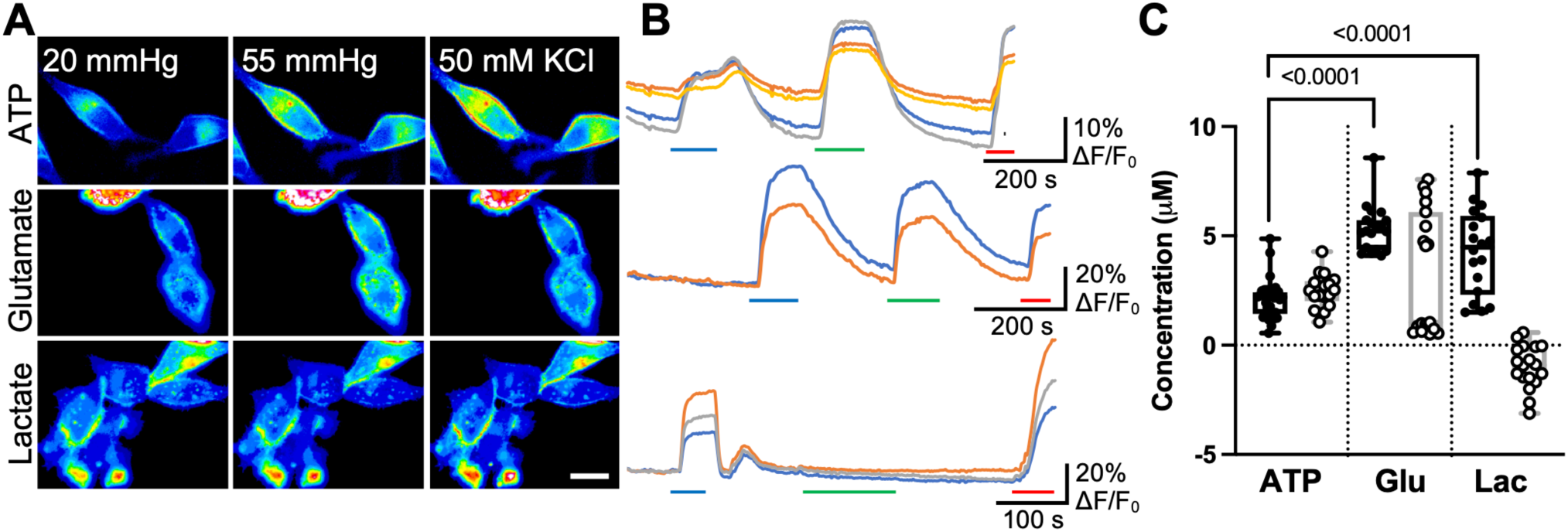
Cx43 hemichannels mediate release of ATP, glutamate and lactate. A, Representative images showing cells from each sensor under each condition. Scale bar represents 20 μm. **B,** Representative traces of normalised fluorescence changes in response to 55 mmHg PCO^2^ (blue bar), 50 mM KCl (green bar) and a 3 μM calibration of the corresponding analyte (red bar). **C,** Summary data showing the median release of analytes through Cx43 hemichannels: ATP (n = 33 (CO^2^), n = 17 (high KCl)), glutamate (n = 19), lactate (n = 18). Data shows CO^2^-dependent release (open circles), and release to a depolarising stimulus (closed circles). Data presented is from at least 3 independent transfections. Kruskal Wallis Anova p<0.0001 for 55 mmHg CO^2^ and p<0.0001 for 50 mM K^+^. Mann Whitney U-tests, p < 0.0001 (CO^2^ ATP vs glutamate), p < 0.0001 (CO^2^ ATP vs lactate). Box and whisker plots with superimposed data points, showing the median (line), the interquartile range (box) and range (whiskers).

While permeability of Cx43 hemichannels to ATP has been elegantly demonstrated (Kang et al., 2008), previous reports suggest that Cx43 hemichannels are apparently not permeable to glutamate or lactate (Hansen et al., 2014; Nielsen et al., 2019). However these previous studies used removal of extracellular Ca^2+^ to unblock the channel, and the channel might thus have a different permeability profile. Whereas zero [Ca^2+^]^ext^ is often achieved by the use of chelators such as EGTA, the genetically encoded fluorescent sensors have some degree of Ca^2+^-dependency. To ensure compatibility with the sensors, we omitted EGTA and Ca^2+^ to lower but not eliminate extracellular Ca^2+^. To ensure this was still sufficient to open the hemichannels, we employed a dye loading assay using the hemichannel-permeable dye, FITC. Under control conditions (PCO^2^ 20 mmHg, 2 mM [Ca^2+^]^ext^) the median pixel intensity was 15.7 (24.72, 14.04). In low [Ca^2+^]^ext^ (PCO^2^ 20 mmHg), the median pixel intensity was 54.3 (86,74, 51.82) (Supplementary Fig. 2). For a comparison to a reliable opening stimulus, we use a PCO^2^ of 55 mmHg to open the hemichannels and permit loading with FITC. This yielded a median pixel intensity of 48.0 (54.83, 40.66) and demonstrates that the low [Ca^2+^]^ext^ solution was an effective stimulus to open Cx43 hemichannels.

Having established the efficacy of low [Ca^2+^]^ext^ at opening Cx43 hemichannels, we assessed the permeation of ATP, glutamate and lactate during this stimulus and compared it to permeation in response to hypercapnia in the same cells. Consistent with our previous findings, hypercapnia evoked the release of ATP, glutamate and lactate. However, low [Ca^2+^]^ext^ significantly reduced the analyte release through Cx43 hemichannels, with a median release of 0.1 (0.86, 0.08) μM ATP, 0.2 (0.76, 0.004) μM glutamate and 0.5 (2.11,-0.25) μM lactate (Supplementary Fig. 3). The nature of the gating mechanism appears to change channel permeability and may account for why our results differ from previous release studies.

Cx31.3 (also called Cx29) is mainly expressed in myelinating cells (Cisterna et al., 2019). There has been some suggestion that Cx31.3 predominantly forms hemichannels and mediates ATP efflux from cells that express this isoform (Sargiannidou et al., 2008; Liang et al., 2011). As Cx31.3 is not CO^2^-sensitive (Butler et al., 2025), we used a depolarising stimulus to open this hemichannel. We were able to demonstrate a median release of 2.6 (2.91, 2.26) μM ATP, 2.3 (2.58, 1.91) μM glutamate and 2.8 (2.92, 2.24) µM lactate (Fig. 5) via Cx31.3 hemichannels. While relatively non-selective, Cx31.3 hemichannels nevertheless show some preference for ATP as the relative permeation of glutamate and lactate is less than that predicted by the electrochemical driving force on these molecules (Tables 1 and 2).

**Figure 5.**
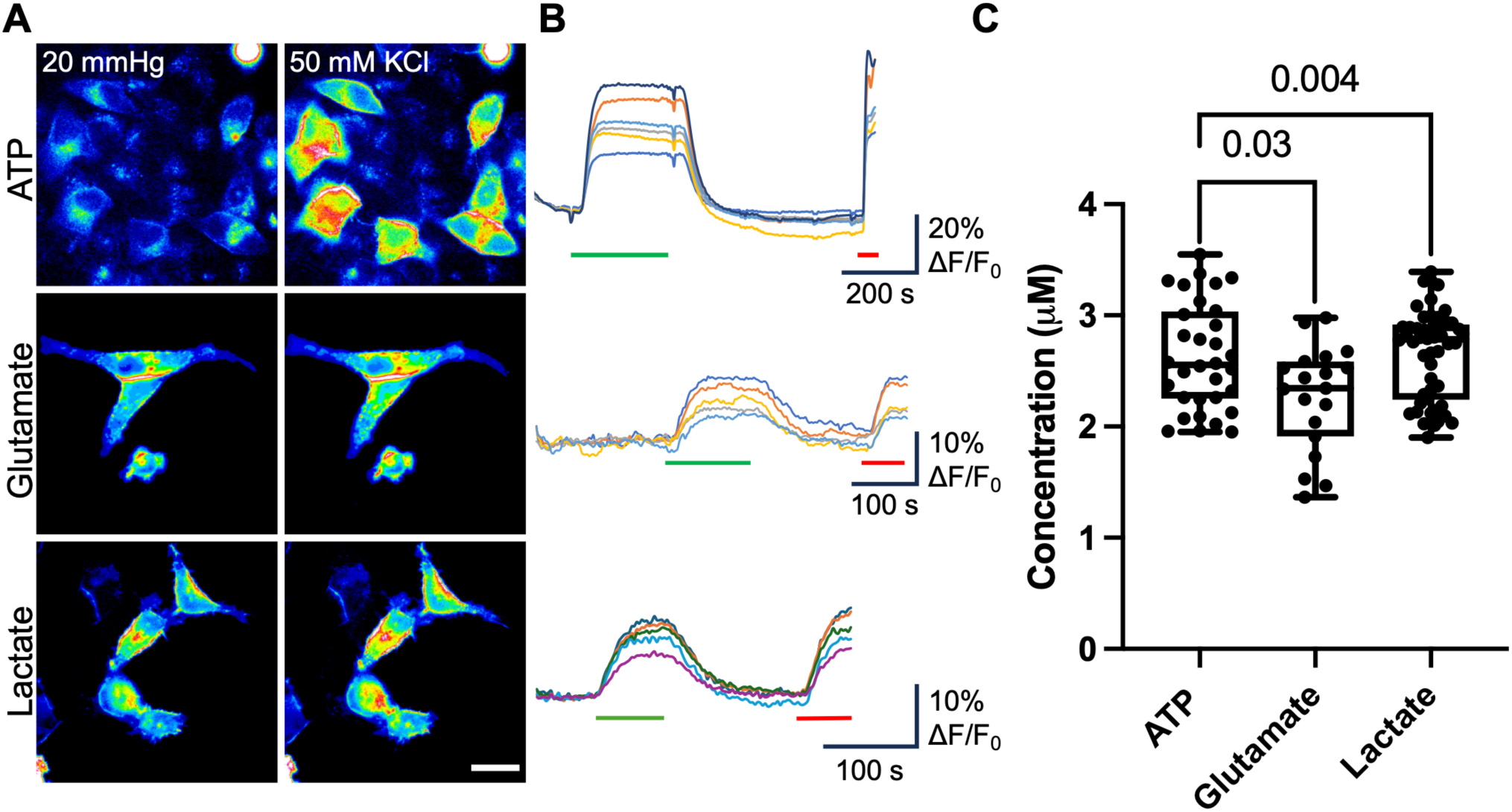
Cx31.3 hemichannels mediates release of ATP, glutamate and lactate. A, Representative images showing cells from each sensor under each condition. Scale bar represents 20 μm. **B,** Representative traces of normalised fluorescence changes in response to 50 mM KCl (green bar), and 3 μM analyte (red bar). **C,** Summary data showing the median ATP (n = 32), glutamate (n = 19) and lactate (n= 50) release from Cx31.3 hemichannels in response to a depolarising stimulus. Data presented is from at least 3 independent transfections. Kruskal Wallis Anova p=0.0174. Mann Whitney *U*-tests, p = 0.004 (ATP vs glutamate) and p=0.03 (lactate vs glutamate). Box and whisker plots with superimposed data points, showing the median (line), the interquartile range (box) and range (whiskers).

### Cx36, Cx46 and Cx50 hemichannels have highly specific permeability profiles

Cx36 acts predominantly as a major neuronal connexin (Condorelli et al., 1998) and forms the gap junctions or electrical synapses that facilitate fast synaptic transmission and synchronous neuronal firing (Srinivas et al., 1999; Deans et al., 2001; Buhl et al., 2003). We assayed the release of ATP, glutamate and lactate from HeLa cells transfected with Cx36. Because Cx36 is insensitive to CO^2^ (Huckstepp et al., 2010a), we used the depolarising stimulus to gate the channel. Surprisingly, we found that of the three analytes, Cx36 was only permeable to ATP (Fig. 6). The median release of ATP evoked by 50 mM KCl was 2.6 (3.27, 2.41) μM, compared to-0.3 (0.22,-0.68) and-0.2 (-0.05,-0.21) μM glutamate and lactate respectively. As the electrochemical driving force for release of glutamate and lactate is about double that of ATP (Tables 1 and 2), and these molecules are smaller than ATP, the differential permeability of Cx36 suggests the existence of a selectivity filter within the pore.

**Figure 6.**
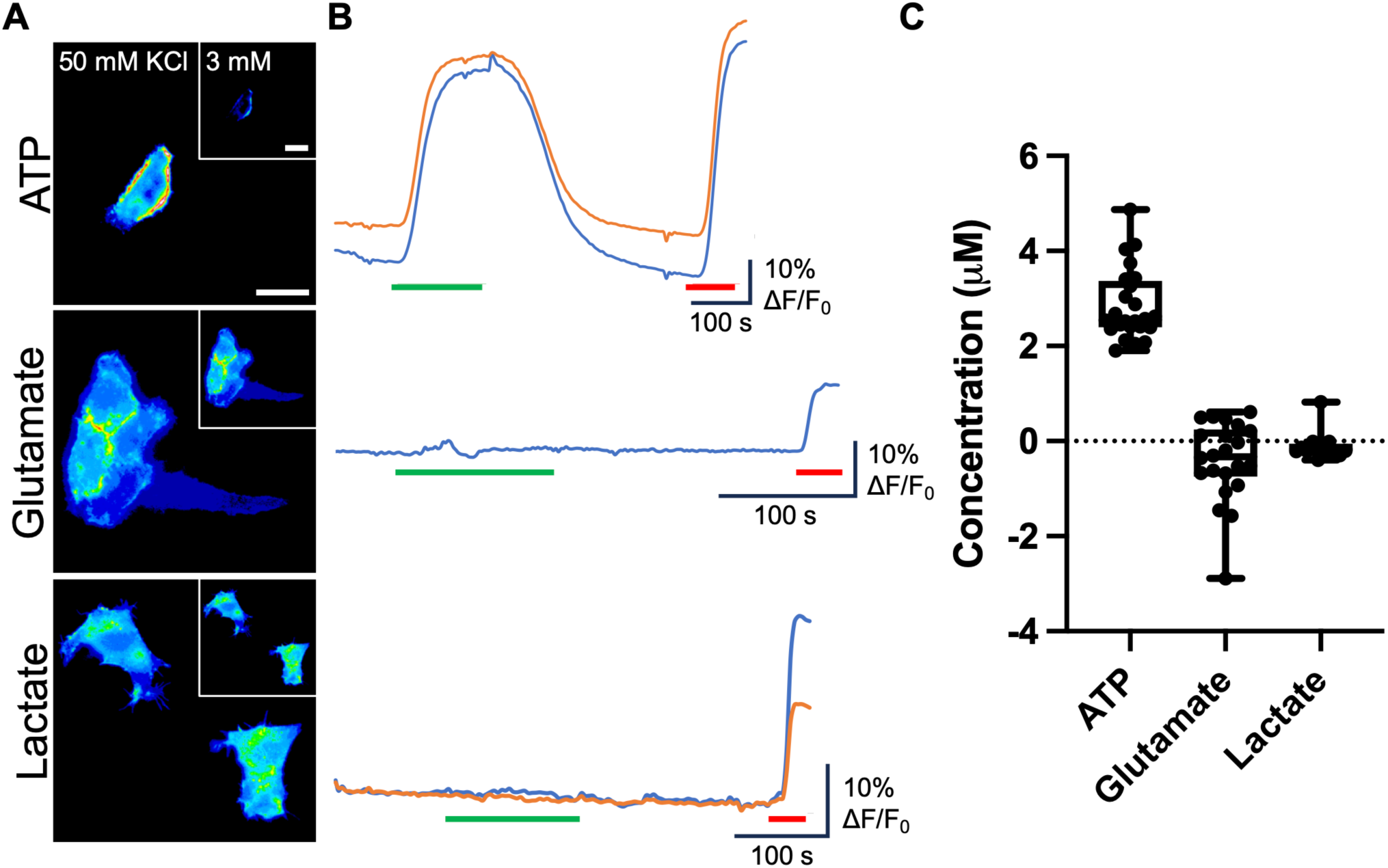
Cx36 hemichannels mediate release of ATP to depolarisation but are impermeable to glutamate and lactate. A, Representative images showing cells from each sensor under each condition: ATP (n = 24), glutamate (n = 22), lactate (n = 15). Inset is 3 mM KCl baseline control. Scale bar represents 20 μm. **B,** Representative traces of normalised fluorescence changes in response to 50 mM KCl (green bar) and 3 μM analyte (red bar). **C,** Summary data depicting ATP release via Cx36 hemichannels and no permeability to glutamate or lactate. Kruskal Wallis Anova p<0.0001. Mann Whitney *U*-tests, p < 0.0001 (ATP vs glutamate, ATP vs lactate). Box and whisker plots with superimposed data points, showing the median (line), the interquartile range (box) and range (whiskers).

Human Cx46 also lacks the carbamylation motif and is not sensitive to CO^2^. We therefore used the high K^+^ stimulus to open Cx46 hemichannels. Like Cx36, Cx46 hemichannels were only permeable to ATP giving a median release of 2.6 μM (2.74, 2.34) (Fig. 7). No glutamate or lactate was released via Cx46 hemichannels (median release-0.07 μM (0.11,-0.28) and 0.2 μM (0.41, - 0.28) for glutamate and lactate respectively, Fig. 7). By contrast, Cx50 hemichannels were readily permeable to glutamate (median release 1.5 μM (1.57, 1.32)) when stimulated by hypercapnia and lactate when stimulated by either hypercapnia or depolarisation (median release 5.8 μM (7.36, 2.55) and 3.4 μM (3.78,-0.55) respectively, Fig 7). However, no ATP could permeate Cx50 hemichannels (Fig. 7).

**Figure 7.**
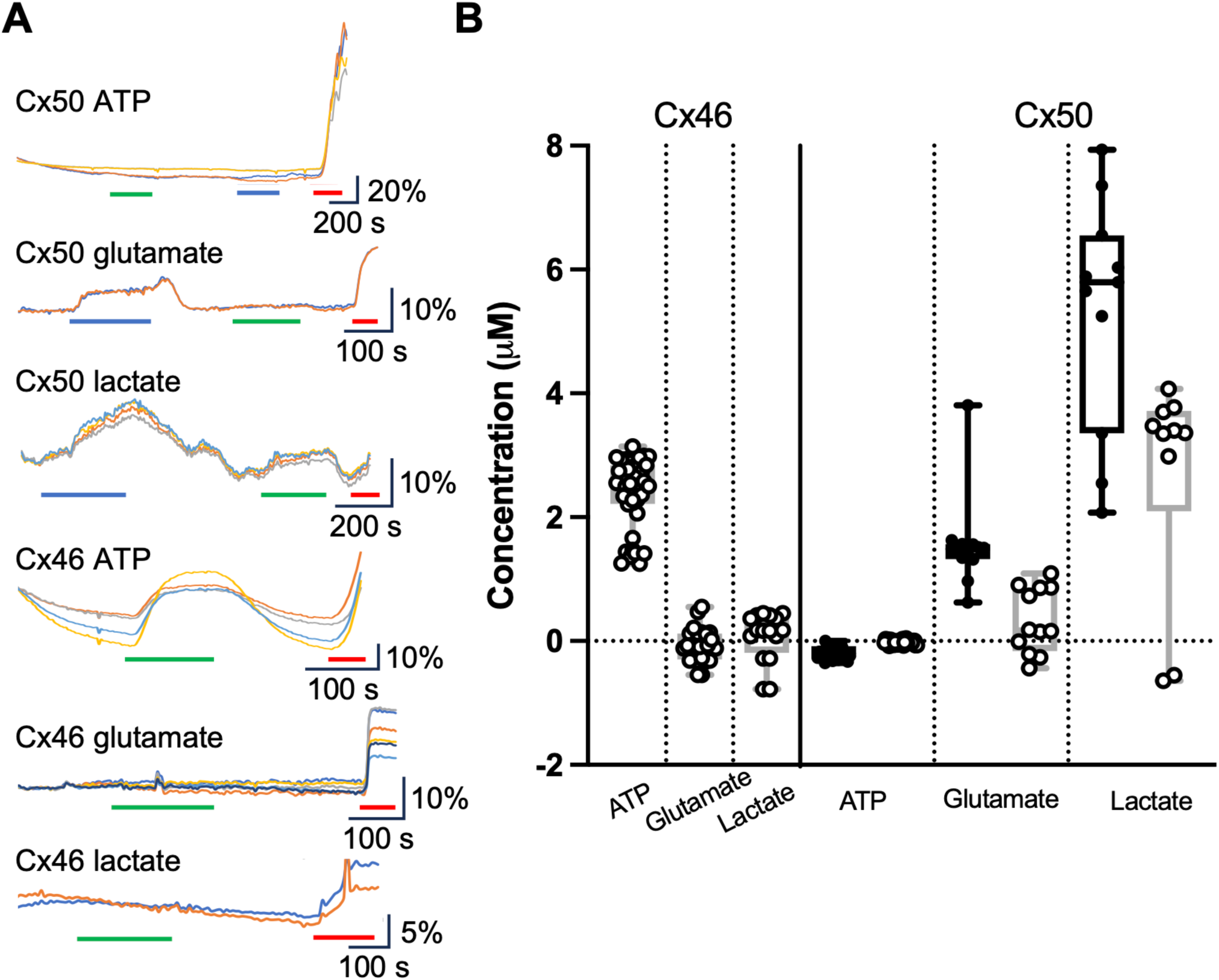
Cx46 and Cx50 hemichannels have opposing permeability profiles. A, Representative traces showing normalised fluorescence changes in response to 55 mmHg (blue bar), 50 mM KCl (green bar) and 3 μM (analyte). **B,** Summary data depicting analyte release under each stimulus: Cx46 ATP (n = 40), Cx46 glutamate (n = 25), Cx46 lactate (n = 16), Cx50 ATP (n = 28), Cx50 glutamate (n = 12), Cx50 lactate (n = 11). Cx50 is CO^2^ sensitive, and thus release is stimulated by 55 mmHg PCO^2^ (open circles), and 50 mM KCl (closed circles). Data presented is from 3 independent transfections. Box and whisker plots with superimposed data points, showing the median (line), the interquartile range (box) and range (whiskers).

### Mutational analysis of the differential permeability of Cx46 and Cx50 hemichannels

Cx46 and Cx50 are structurally quite similar (Myers et al., 2018; Flores et al., 2020; Yue et al., 2021) yet their hemichannels have markedly different permeability profiles. These two connexins therefore offer an opportunity to explore the mechanistic basis of differential hemichannel permeability. The N-termini fold into the gap junction pore to form the narrowest point, with hydrophobic residues anchoring the helix to transmembrane regions 1 and 2 (TM1/2) to stabilise the open state (Myers et al., 2018). Aligning the N-termini sequences of Cx46 and Cx50 shows that they have a difference in the overall net charge (respectively 0 and-2 for Cx46 and Cx50, Fig. 8). The first divergence is at position 9, where Cx46 has a positively charged arginine and Cx50 has an asparagine. At position 13, Cx50 has a negatively charged glutamate residue where Cx46 has a neutrally charged asparagine.

**Figure 8.**
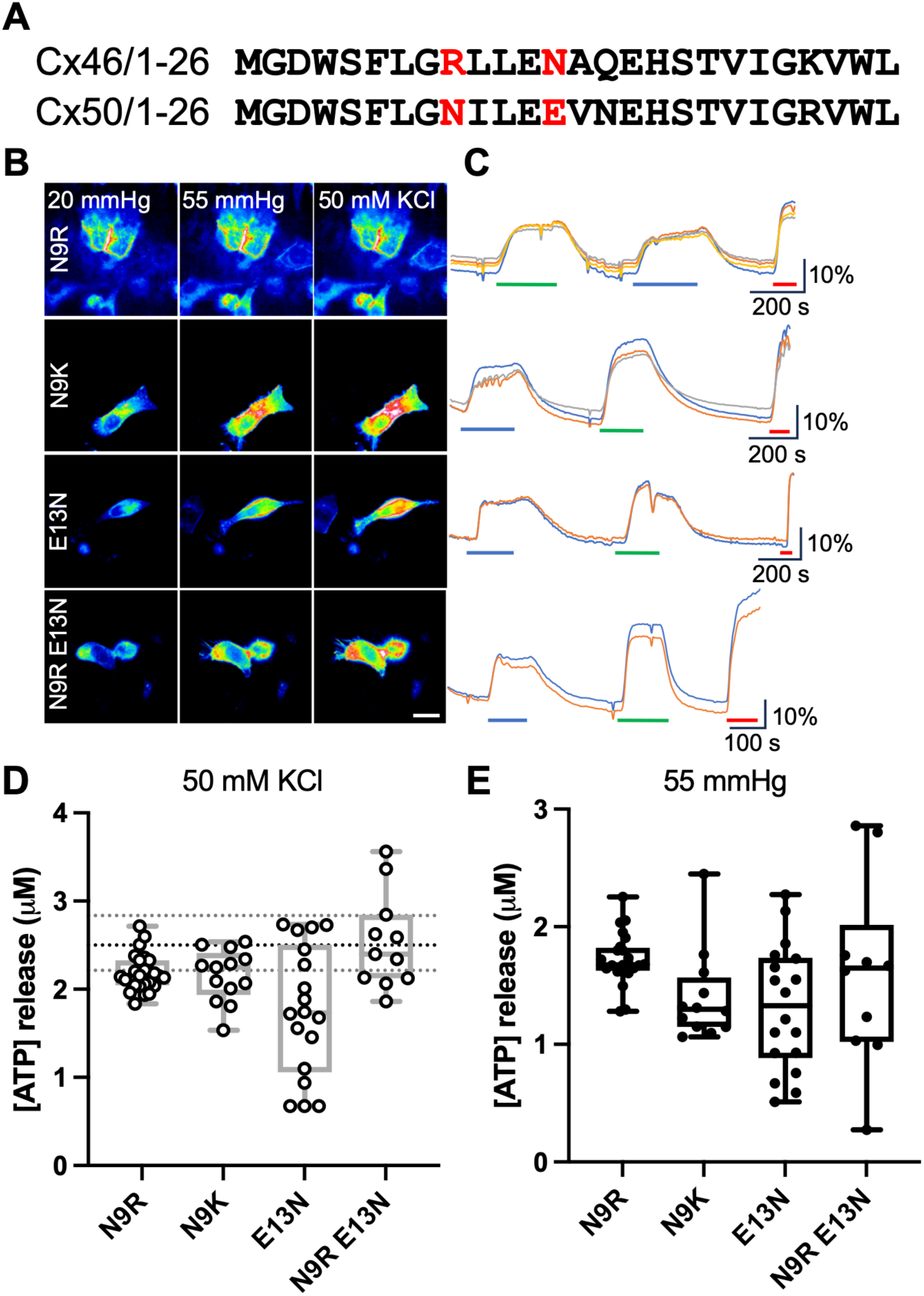
N-terminal mutations make Cx50 hemichannels permeable to ATP. A, Sequence alignments for residues 1-26 of Cx46 and Cx50, to show the residues in Cx50 that have been mutated to those in Cx46 (red letters). **B,** Representative images showing cells from each sensor under each condition. Scale bar represents 20 μm. **C,** Representative traces of normalised fluorescence changes in response to 55 mmHg (blue bar), 50 mM KCl (green bar) and 3 μM ATP (red bar). **D,** Summary data showing ATP release in response to depolarisation from Cx50 mutant hemichannels: N9R (n = 26), N9K (n = 13), E13N (n = 18), N9R E13N (n = 11). Data shows CO^2^-dependent release (open circles), and release to a depolarising stimulus (closed circles). The dotted lines show the median and 95% confidence limits for ATP release from Cx46 in response to depolarisation. Kruskal Wallis Anova (all 4 mutants plus Cx46^WT^) p = 0.0009. Mann Whitney *U*-tests, p = 0.0018, p = 0.0155 and p = 0.0004 (N9R, N9K and E13N respectively, where release evoked by 50 mM KCl is compared to Cx46^WT^) and p = 0.96 (Cx46^WT^ vs Cx50^N9R E13N^). Box and whisker plots with superimposed data points, showing the median (line), the interquartile range (box) and range (whiskers). **E,** Summary data demonstrating CO^2^-evoked ATP release form the Cx50 mutant hemichannels N9R (n = 26), N9K (n = 13), E13N (n = 18), N9R E13N (n = 11). Box and whisker plots with superimposed data points, showing the median (line), the interquartile range (box) and range (whiskers).

To explore the possible roles of the difference in net charge, we introduced the single mutations N9R, E13N and N9K into Cx50 to make it more like Cx46. These mutations gave a gain of ATP permeability to Cx50 hemichannels, though this was still significantly less than that of Cx46^WT^ hemichannels (Fig. 8). The double mutation, Cx50^N9R E13N^, gave an increase in ATP release that matched that of Cx46^WT^ hemichannel release (for a depolarising stimulus), with a median release of 2.4 (3.37, 2.07) μM from the mutant connexin compared to 2.5 (2.74, 2.34) for Cx46^WT^ hemichannels. We have therefore shown that the net charge of the N-terminus, and the specific residues N9 and E13, act to regulate the ATP permeability of Cx50 hemichannels.

We next examined whether, if we made the N-terminus of Cx46 more like that of Cx50, we could change the permeability profile of Cx46 hemichannels to match those of Cx50. We therefore made the double mutation R9N,N13E in Cx46. However, this did not diminish ATP permeation via Cx46 hemichannels (Fig. 9). The median release from Cx46^R9N,N13E^ hemichannels was 2.8 (3.10,2.47), compared with 2.5 (2.74, 2.34) μM for Cx46^WT^ hemichannels. This indicates that the regulation of permeability of Cx46 hemichannels to ATP is more complex than just the net charge of the N-terminus.

**Figure 9.**
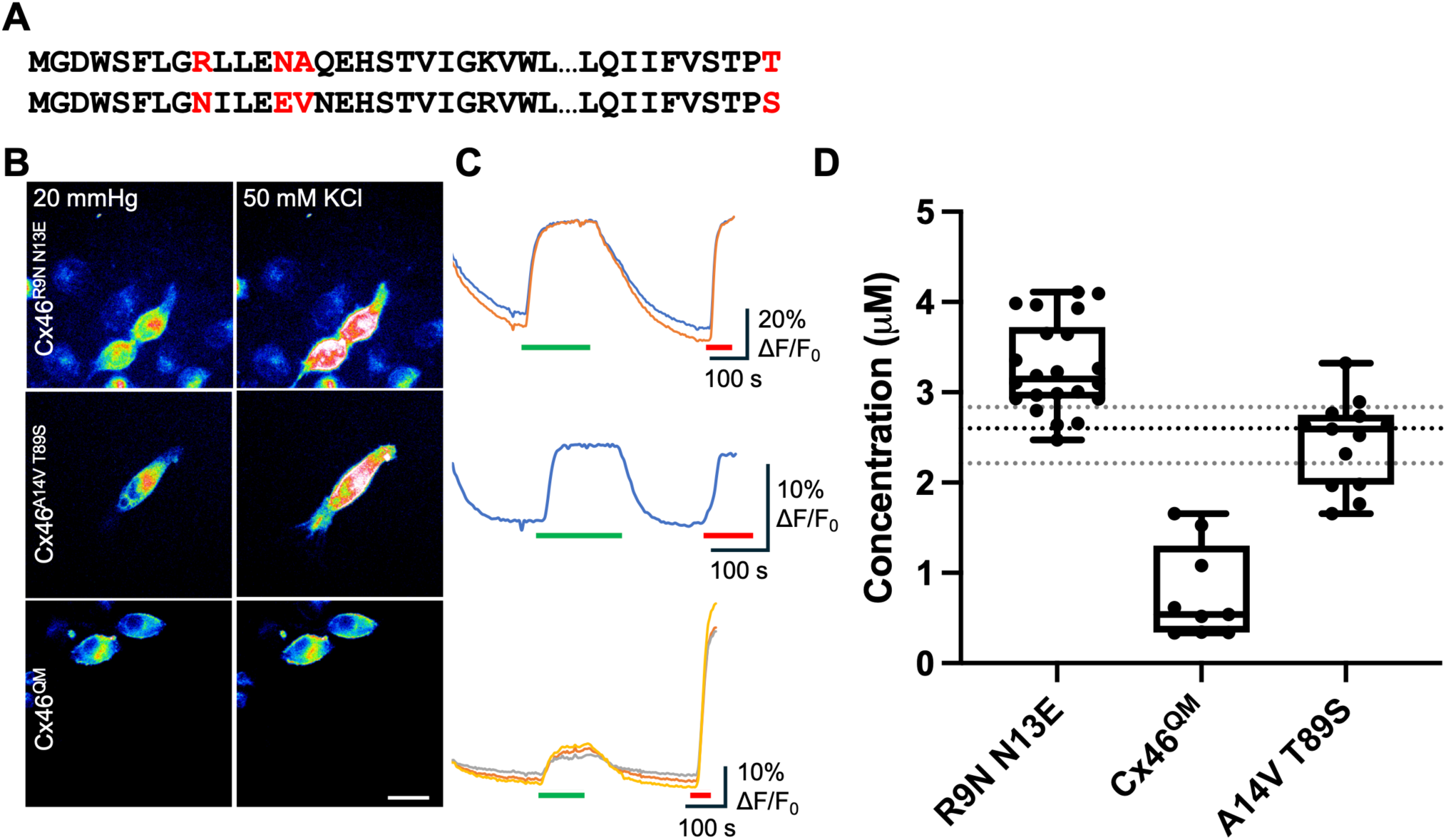
The permeability of Cx46 hemichannels to ATP appears to be reliant on N-terminal charge and interactions between the N-terminus and TM2. A, Sequence alignments from 1-26 and 80-89 for Cx46 (top) and Cx50 (bottom), to show the residues in Cx46 which have been mutated to correspond to those in Cx50 (red). **B,** Representative images showing cells expressing Cx46^R9N N13E^, Cx46^A14V T89S^ and Cx46^QM^ under each condition. Scale bar represents 20 μm. **C,** Representative traces showing normalised change in fluorescence to 50 mM KCl (green bar), and 3μM ATP (red bar). **D,** Summary data depicting the median ATP release from hemichannels of Cx46^WT^ (the dotted lines show the median and 95% confidence interval for Cx46^WT^), Cx46^R9N N13E^ (n = 22), Cx46^QM^ (n = 9), and Cx46^A14V T89S^ (n = 14). Mann Whitney *U*-test, p < 0.0001 (Cx46^WT^ vs Cx46^QM^). Box and whisker plots with superimposed data points, showing the median (line), the interquartile range (box) and range (whiskers).

We considered the possibility that interactions between the N-terminus and TM2 may differ between Cx46 and Cx50 and this could permit ATP permeation even if the net charge of the N-termini were negative. Both Cx46 and Cx50 have hydrophobic residues at position 14: Cx50 has a valine, but Cx46 has an alanine. We therefore used experimentally determined structures of these connexins (Jaradat et al., 2022) to identify possible interacting residues in TM2. This highlighted residue 89 as potentially important: in Cx50 this is serine, but in Cx46 it is threonine. We sought to make these interactions in Cx46 more similar to those occurring in Cx50 by introducing the double mutation A14V,T89S. We note that simple introduction of the Cx50 N-terminal helix into Cx46 has been reported as resulting in non-functional gap junction channels due to a steric clash between V14 and T89 in the chimaeric channel (Yue et al., 2021). By making two mutations in Cx46, A14V and T89S, we have avoided this clash. The double mutation A14V, T89S did not by itself alter ATP permeation (Fig. 9), indicating that the Cx46^A14V,T89S^ hemichannel gated normally to voltage. However, when A14V and T89S were then combined with R9N and N13E, the quadruply mutated Cx46^R9N,N13E,A14V,T89S^ hemichannels (Cx46^QM^) exhibited significantly reduced ATP permeation compared to the wild-type hemichannels, with a median release of 0.54 (1.53, 0.34) μM (Fig. 9).

To check that the mutations of Cx46 and Cx50 did not alter the trafficking and membrane localisation of the resulting hemichannels, we performed confocal analysis to quantify the colocalization of the mCherry tag with a plasma membrane stain DiO. Using Manders analysis, we found that the proportion of mCherry colocalised with DiO was the same in all mutations and their respective wildtype connexins (Fig. 10 and Supplementary Figure 4).

**Figure 10.**
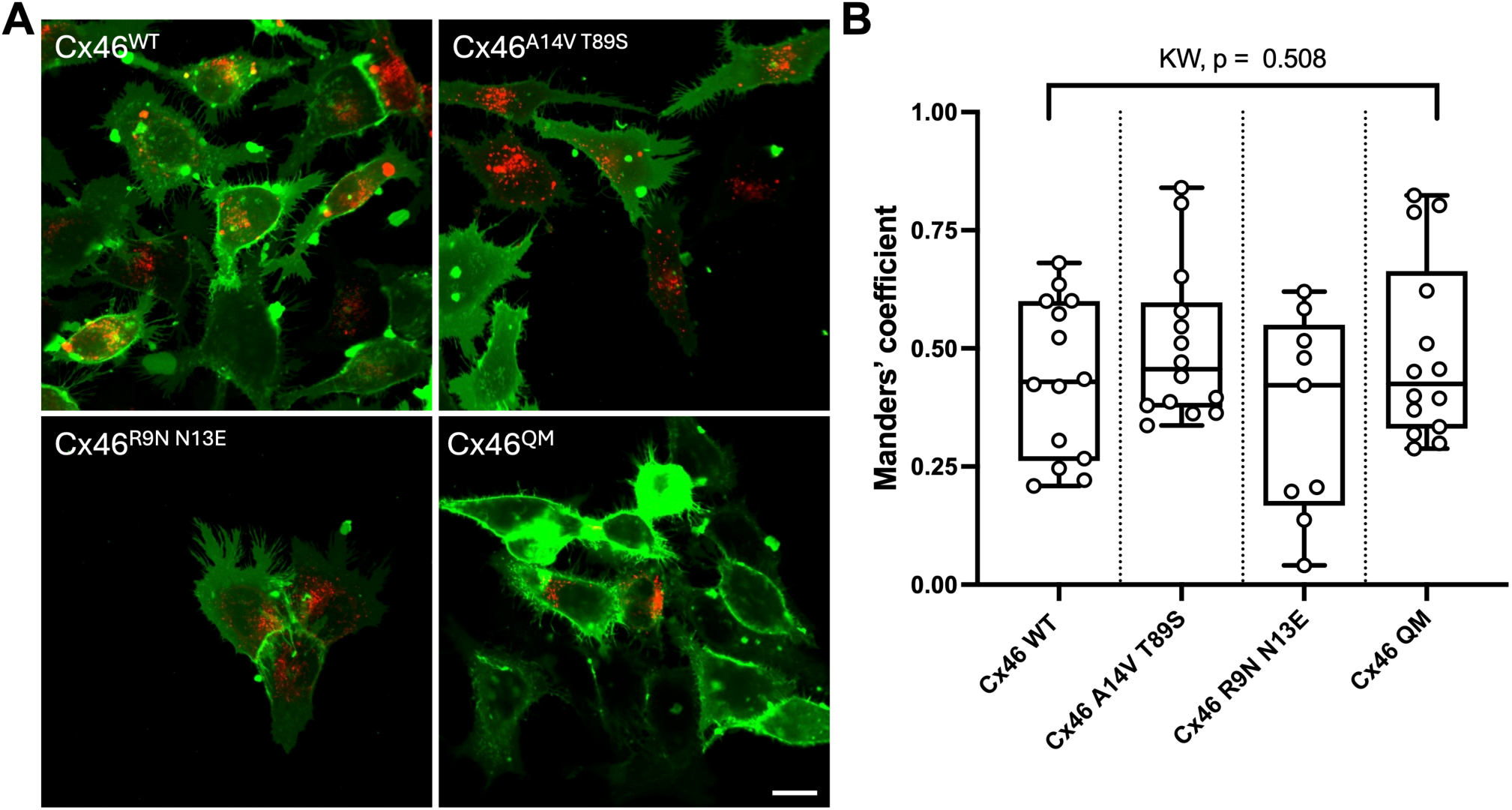
The plasma membrane distribution of Cx46 is unaltered by mutations that alter its permeability to ATP. A, Confocal images of DiO staining (green) and the mCherry-tagged Cx46 expression (single optical plane). Scale bar 15 µm. **B,** Manders’ analysis of colocalization shows that the proportion of mCherry colocalised with DiO is the same for Cx46^WT^, and the three mutated variants. Data is from 3 independent transfections. Kruskal Wallis Anova, p=0.508.

One possible interpretation of the reduced permeability to ATP of Cx46^QM^ is that the quadruple mutation simply reduces overall hemichannel permeability rather than having a specific effect on that of ATP. To evaluate this, we examined depolarisation dependent loading of FITC into HeLa cells expressing either Cx46^WT^ or Cx46^QM^ (Fig. 11 A,B). We found that FITC still permeated into cells expressing Cx46^QM^ and was not significantly different from the permeation observed in those expressing Cx46^WT^. This suggests that the quadruple mutant has a somewhat selective effect on ATP permeation. We also checked whether Cx46^QM^ might display enhanced glutamate permeability (as per Cx50 hemichannels). However, this was not the case (Fig. 11 C,D) suggesting that the determinants of permeation of these two analytes in Cx46 are somewhat independent.

**Figure 11.**
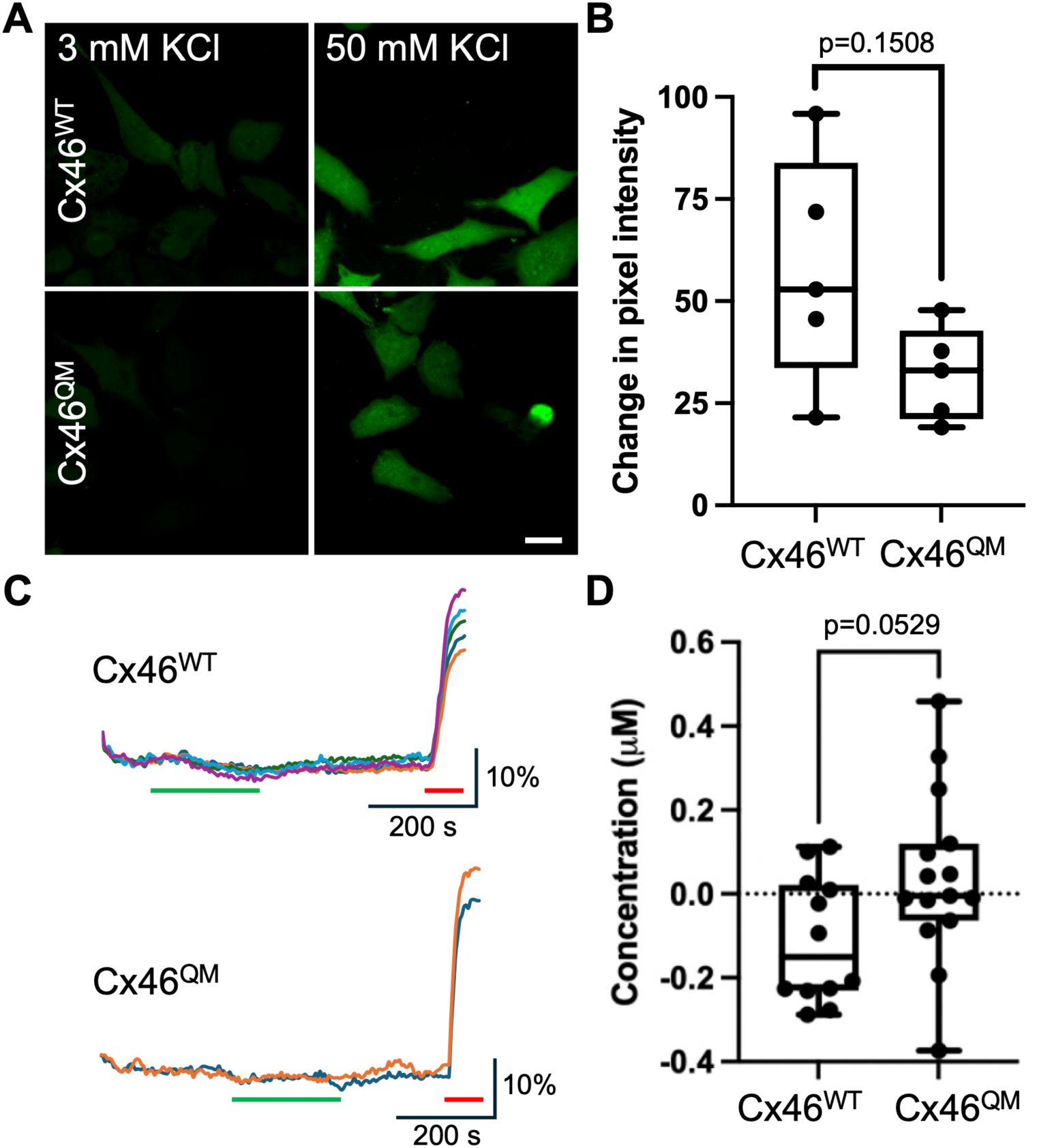
Cx46^QM^ hemichannels remain permeable to FITC and do not gain permeability to glutamate. A, Representative images showing cells expressing Cx46^WT^ or Cx46^QM^ loaded with FITC under each condition. Scale bar represents 15 μm. **B,** Summary data depicting the median change in fluorescence pixel intensity between 3 mM KCl and 50 mM KCl for cells expressing Cx46^WT^ or Cx46^QM^ from 5 independent transfections. There is no significant difference in FITC loading into the Cx46^QM^ compared to Cx46^WT^ (Mann Whitney *U*-Test). For Cx46^WT^, 43 cells in 20 mmHg and 35 cells in 50 mM KCl were analyzed. For Cx46^QM^, 79 cells in 20 mmHg and 79 cells for Cx46^QM^ in 50 mM KCl were analyzed. Box and whisker plots with superimposed data points, showing the median (line), the interquartile range (box) and range (whiskers). **C,** iGluSnFR traces for Cx46^WT^ and Cx46^QM^ showing that 50 mM KCl (green bar) does not evoked glutamate release. Red bar 3 µM Glu. **D,** Summary data showing median glutamate release from Cx46^WT^ (n=12) and Cx46^QM^ (n=15) (3 independent transfections, Mann Whitney *U*-test). Box and whisker plots with superimposed data points, showing the median (line), the interquartile range (box) and range (whiskers).

## Discussion

This study explored whether there is selectivity to the release of small molecules from connexin hemichannels expressed in HeLa cells. To assess their relative permeability via the different connexins, we need to understand the electrochemical driving force on the three metabolites. Whereas this driving force on each metabolite is likely to be cell-type specific, the permeability properties of a connexin hemichannel pore are encoded by the protein and are likely to remain invariant between different cell types. As ATP, glutamate and lactate are charged, we can use the Nernst equation, along with typical concentrations of these molecules in HeLa cells (Piva and McEvoy-Bowe, 1998; Imamura et al., 2009; San Martin et al., 2013) to calculate the equilibrium potential (Table 1). If we assume that the concentrations of these metabolites are scattered around a mean value that is consistent across all HeLa cells, sufficient recordings of release should statistically reflect these transmembrane concentrations. Based on this analysis, if release through connexin hemichannels were to be non-selective, the relative proportions of release should correspond to the relative electrochemical driving forces. Thus, for a completely non-selective connexin hemichannels we would expect to see ATP, glutamate and lactate released approximately in the proportion 1:1.8:1.4 (assuming a resting potential of - 60 mV) to hypercapnia and in the proportion 1:2.1:1.6, for the depolarising stimulus. Any notable deviation from these proportions would indicate that there is some selectivity to release of small molecules through connexin hemichannels (Table 2).

Our study is the first to produce a comprehensive permeability profile of a wide range of connexin isoforms for release of physiological metabolites with physiological opening stimuli. Connexin hemichannels fall into two broad categories: relatively non-selective (Cx26, Cx32, Cx43, Cx31.3); and highly selective (Cx36, Cx46 and Cx50). For Cx26 and Cx32 hemichannels, the release of glutamate and lactate relative to ATP is more than predicted by the electrochemical driving force, suggesting that the smaller molecules may permeate more readily than ATP (Table 2). Cx32 hemichannels in particular show enhanced permeability to lactate (nearly 4 times that predicted by driving force). Interestingly, when Cx32 hemichannels were opened by depolarisation, the relative release of analytes followed that predicted by the electrochemical driving force more closely. For Cx43 hemichannels, when opened by hypercapnia, the relative release of ATP, glutamate and lactate follows very closely the pattern predicted by the electrochemical driving force (Table 2). Cx31.3 hemichannels show some preference for ATP over glutamate and lactate but is nevertheless permeable to these smaller molecules (Table 2).

Rather surprisingly, Cx50 hemichannels are impermeable to ATP. To our knowledge this is the only connexin hemichannel that ATP cannot permeate. Traditionally, connexin hemichannel permeability studies have studied selectivity by increasing the size of fluorescent dyes (Weber et al., 2004; Harris, 2007), with the idea being that any molecule that was below the limiting pore diameter (around 12 Å) should permeate. This would suggest minimal selectivity between permeants, leaving the major driving force as intracellular concentration. Our results modify this idea. Although the smaller molecules glutamate and lactate do permeate the relatively non-selective channels more easily, lactate being the smallest molecule should permeate all connexin hemichannels: but it cannot permeate Cx36 or Cx46 hemichannels, whereas ATP, a much larger molecule can.

We also find that the permeability profile of the hemichannel can alter with the nature of the gating stimulus. For those connexins that are directly CO^2^ sensitive, opening the hemichannel by hypercapnia seems to give greater release than depolarisation (summarized in Table 2). One possibility is that 50 mM KCl, while predicted to depolarise the cell by about 70 mV, may not sufficiently depolarise the membrane to obtain full channel opening. We also find that lowered [Ca^2+^]^ext^ seems to be the least effective stimulus for release. While this manipulation may unblock the channel (by removing the ring of bound Ca^2+^ ions), the N-termini may still partially block the channel and alter the permeation pathway in a way that may not happen with a more physiological stimulus.

### The selective permeability of connexins may match their physiological roles

That there are 21 connexin genes in the human genome, indicates distinct fundamental roles in physiological processes. The need for so many isoforms suggests functional specialisation - combinations of properties that match connexin to function. Sensitivity to gating stimuli, and permeability to small molecules are two important properties that may determine which functional roles connexins are suited to.

The relatively non-selective connexins, Cx26, Cx32 and Cx43, have different CO^2^ sensitivity profiles. Cx26 is suited to detection of systemic PCO^2^ and has a role in the control of breathing (Huckstepp et al., 2010b; van de Wiel et al., 2020; Dale, 2021). Cx32 requires much higher levels of PCO^2^ to open and may be more suited to detecting local CO^2^ production (Huckstepp et al., 2010a; Dospinescu et al., 2019; Butler and Dale, 2023). It is interesting that Cx32 hemichannels are highly permeable to lactate when opened by hypercapnia. This lends support to an attractive hypothesis that Cx32 hemichannels may detect hotspots of metabolic activity and, by opening and permitting release of lactate, could provide metabolic support for highly active cells (Barros, 2013). Cx43 with its extensive C-terminus interacts with many other proteins (Iacobas et al., 2003; Iacobas et al., 2007), yet is also CO^2^ sensitive (Dospinsecu et al., 2025) and its hemichannels are partially open under physiological conditions (Chever et al., 2014; Turovsky et al., 2020).

We also discovered a group of connexins with highly selective hemichannel permeability profiles: Cx36, Cx46 and Cx50. Cx36 is expressed predominantly in neurons and the hemichannel is impermeable to glutamate and lactate. The lack of glutamate permeability may be functionally significant as most excitatory neurons use this as their main neurotransmitter. This lack of permeability may ensure that glutamate release is tightly regulated under physiological conditions and occurs mainly via vesicular exocytosis at synaptic sites. As lactate is an effective metabolite for oxidative phosphorylation in neurons, the lack of permeability of Cx36 hemichannels to lactate would prevent unregulated efflux of lactate from neurons.

Cx46 and Cx50 are expressed almost exclusively in the lens of the eye (Mathias et al., 2010; Berthoud and Ngezahayo, 2017). Cx46 and Cx50 form gap junctions between the lens epithelial and fibre cells, and between lens fibre cells. Cx50 is also present as hemichannels in lens fibre cells. As lens fibre cells mature, they lose their intracellular organelles including mitochondria (Bassnett, 2002) and thus the ability to make ATP via oxidative phosphorylation. One can speculate that having ATP permeable Cx46 gap junction channels may be valuable in allowing diffusion of this key metabolite from the metabolically active cells in the lens into the relatively inactive lens fibre cells. Equally, the lack of ATP permeability of Cx50 hemichannels may permit preservation of this scarce resource in the lens fibre cells.

### The N-terminus is a fundamental determinant of selectivity

Considerable evidence suggest that the N-terminus is part of the gating mechanism of connexins. Depending on the isoform, the N-terminus projects into the pore to form an occluding plug or forms a cap at the cytoplasmic vestibule also to close it. In the open state the N-termini line the pore and thus experience the membrane electric field. This may explain why increasing the charge of the Cx50 N-terminus appears to enhance the voltage-gating of the mutants (Figs 7 & 8) (Kalmatsky et al., 2009). As the N-terminus lines the entrance to the pore (Maeda et al., 2009; Myers et al., 2018; Flores et al., 2020; Yue et al., 2021; Brotherton et al., 2022; Jaradat et al., 2022; Brotherton et al., 2024) it may regulate selectivity to small molecules. Our evidence suggests that this is the case: by manipulating the charge on the N-terminus of Cx50 to make it more like Cx46, we were able to create a mutant hemichannel that gained ATP sensitivity over the wild type Cx50 hemichannel. We note that the mutation Cx50^N9R^ increased the permeability to ATP through the hemichannel, despite this mutation being reported to decrease the conductance of the Cx50 gap junction channel (Yue et al., 2021). However, the reverse set of mutations-changing charge on the N-terminus of Cx46 to make it more Cx50-like did not diminish ATP permeability of the mutant Cx46 hemichannel. This shows that other mechanisms can be important.

Our data suggests that residues that alter the interaction between the N-terminus and the lining of the pore-notably TM2 – play a role in hemichannel selectivity (Brotherton et al., 2022; Brotherton et al., 2024). We identified Ala14 and Thr89 of Cx46 as residues that might be important in this interaction and mutated those to their equivalent in Cx50 (A14V,T89S). Thr89 in Cx46 is equivalent to Ala88 of Cx26, which interacts with Val13 on the N terminus (Brotherton et al., 2022). Substitution of larger residues (A88V) change Cx26 channel function and are pathological (Koval, 2013; Meigh et al., 2014). However, the double mutation in which a smaller residue is substituted (T89S) avoids the steric clash between V14 and T89 previously reported in Cx46-50 chimaeric channels which were non-functional (Yue et al., 2021). The double mutations A14V and T89S or R9N and N13E by themselves did not alter ATP permeability. However, when all four mutations were combined to create Cx46^QM^ there was a considerable reduction of ATP release compared to Cx46^WT^. The effect of the quadruple mutation FITC permeation was not statistically significant. This suggests that rather than simply resulting in a poorly permeable channel, the quadruple mutation has a somewhat selective effect on the ATP permeation pathway. Our study not only highlights the importance of the N-terminus in determining selectivity of connexin hemichannels to small molecules, but also that even in two very closely related connexins there are other structural elements that control permselectivity.

## Materials and Methods

### Connexin mutagenesis

cDNAs for the Cx26 and Cx32 genes were synthesised by Genscript and for the Cx46, Cx50, Cx36, Cx43 and Cx31.3 genes by IDT. These were subsequently subcloned into pCAG-GS-mCherry vector prior to transfection. Point mutations were introduced using Gibson assembly. Overlapping fragments both containing the desired mutation were PCR amplified with primers (IDT). Successful mutagenesis was confirmed using Sanger sequencing (GATC Biotech). Double mutants (Cx50 ^N9R E13N^, Cx46^R9N N13E^) were cloned using successive Gibson assemblies. All Cx constructs were inserted upstream of an mCherry tag, linked via a 12 AA linker (GVPRARDPPVAT).

### Cell culture

We used HeLa DH cells as an expression system as they exhibit very low expression of endogenous connexins and have been used for this purpose by numerous authors over a long period (Elfgang et al., 1995; Choi et al., 2020). Parental HeLa DH cells (ECACC 96112022 RRID:CVCL_2483) were cultured with low-glucose DMEM (Merck Life Sciences UK Ltd, CAT# D6046) supplemented with 10% foetal bovine serum (Labtech.com, CAT# FCS-SA) and 5% penicillin/streptomycin. The HeLa cells were free from mycoplasma. PCR of cell culture supernatant was regularly performed to check for mycoplasma contamination using the EZ-PCR mycoplasma detection kit (SARTORIUS, 20-700-20). Cells were seeded onto coverslips at a density of 4×10^4^ cells per well. Cells were transiently transfected to co-express one connexin isoform and one genetically encoded fluorescent sensor: pDisplay-GRAB_ATP1.0-IRES-mCherry-CAAX was a gift from Yulong Li (Addgene plasmid # 167582; http://n2t.net/addgene:167582; RRID:Addgene_167582) (Wu et al., 2022).

pAEMXT-eLACCO1.1 was a gift from Robert Campbell (Addgene plasmid # 167946; http://n2t.net/addgene:167946; RRID:Addgene_167946) (Nasu et al., 2021). To improve expression of eLACCO1.1, this construct was subcloned into the iGluSnFR exxpression vector backbone. Sequences were verified with Sanger sequencing (GATC).

pCMV(MinDis).iGluSnFR was a gift from Loren Looger (Addgene plasmid # 41732; http://n2t.net/addgene:41732; RRID:Addgene_41732) (Marvin et al., 2013).

A mixture of 1μg of DNA from pCAG-Cx-mCherry construct and 1μg sensor with 3μg PEI was added to cells for 4-8h. Cells were imaged 48 hours after transfection.

### aCSF

Control (20 mmHg PCO^2^) – 140 mM NaCl, 10 mM NaHCO^3^, 1.25 mM NaH2PO^4^, 3mM KCl, 1 mM MgSO4.

Control (35 mmHg PCO^2^) – 124 mM NaCl, 26 mM NaHCO^3^, 1.25 mM NaH^2^PO^4^, 3mM KCl, 1 mM MgSO^4^.

Hypercapnic (55 mmHg PCO^2^) - 100 mM NaCl, 50 mM NaHCO^3^, 1.25 mM NaH^2^PO^4^, 3 mM KCl, 1 mM MgSO^4^.

Hypercapnic (70 mmHg PCO^2^) - 70 mM NaCl, 80 mM NaHCO^3^, 1.25 mM NaH^2^PO^4^, 3 mM KCl, 1 mM MgSO^4^.

High K^+^ (20 mmHg) – 93 mM NaCl, 10 mM NaHCO^3^, 1.25 mM NaH^2^PO^4^, 50 mM KCl, 1 mM MgSO^4^.

For high K^+^ in 35 mmHg, the recipe is the same but with 77 mM NaCl.

All solutions had 10 mM D-glucose and 2 mM CaCl^2^ (or MgCl^2^ when [Ca^2+^]^0^ solution was desired) added just before use and saturated with 98% O^2^/2% CO^2^ (20 mmHg), 95% O^2^/5% CO^2^ (carbogen) (35 mmHg), or carbogen plus CO^2^ (55 mmHg and 70 mmHg). CO^2^ in all solutions was adjusted to give a pH of ∼7.4.

### Live cell fluorescence imaging and analysis

Cells were transiently transfected with one pCAG-Cx-mCherry construct and one genetically encoded fluorescent sensor 48 hours prior to imaging. Cells were perfused with control aCSF until a stable baseline was reached, before perfusion with either hypercapnic or high K^+^ aCSF. Once a stable baseline was reached after solution change, cells were again perfused with control aCSF and when a stable baseline reached, recordings were calibrated by direct application of 3 μM of the corresponding analyte.

All cells were imaged by epifluorescence (Scientifica Slice Scope, Cairn Research OptoLED illumination, 60x water Olympus immersion objective, NA 1.0, Hamamatsu ImagEM EM-SSC camera, Metafluor software). cpGFP in the sensors were excited by a 470 nm LED, with emission captured between 504-543 nm. Connexin constructs have a C-terminal mCherry tag, which is excited by a 535 nm LED and emission captured between 570-640 nm. Only cells expressing both cpGFP and mCherry were selected for recording, with cpGFP images acquired every 4 seconds. For each condition, at least 3 independent transfections were performed with at least 2 coverslips per transfection.

Analysis of all experiments was carried out in ImageJ. Images were opened as a stack and stabilised (Li, 2008). ROIs were drawn around cells co-expressing both sensor and connexin. Median pixel intensity was plotted as normalised fluorescence change (ΔF/F^0^) versus time to give traces of fluorescence change. The amount of analyte release was quantified as concentration by normalising to the ΔF/F^0^ caused by application of 3 μM of analyte, which was within the linear portion of the dose response curve for each sensor. Release froma single cell was considered to be a statistical replicate.

### Confocal imaging of membrane localisation

HeLa cells were transfected with mCherry-tagged wild type and mutant versions of Cx46 and Cx50. After 48h, the cells were washed 3 times with PBS, fixed with 4% paraformaldehyde in PBS for 30 mins and then washed 3 times with PBS. Cells were then incubated with serum free DMEM containing 2.5 μM DiO (3,3′-Dioctadecyloxacarbocyanine perchlorate, Sigma-Aldrich Cat#D4292) for 15 minutes. After three washes in PBS, coverslips were mounted inverted on glass microscope slides in a Fluorshield™ with DAPI mounting medium (Sigma-Aldrich, Cat# F6057). Slides were sealed and imaged on a Zeiss 880 LSM confocal microscope using the 488 and 561nm excitation wavelengths for DiO and mCherry respectively.

Colocalization analysis between the mCherry tag of the connexin variants and the DiO membrane stain was performed with Fiji and the JaCoP plugin (Bolte and Cordelieres, 2006). ROIs were drawn around mCherry positive cells and the image surrounding the ROI was removed. The Manders’ co-efficient (Manders et al., 1993) was used as a quantitative assessment of colocalisation of mCherry with DiO, and hence the membrane localisation of the connexin constructs. Thresholds were set to a limit that included DiO membrane staining but excluded any diffuse background fluorescence.

### Dye loading

Cells were transiently transfected with pCAG-Cx43-mCherry, pCAG-Cx46^WT^-mCherry or pCAG-Cx43^QM^-mCherry constructs 48 hours prior to experiments. After washing in 20 mmHg aCSF, cells were perfused with one of the following solutions: 20 mmHg aCSF, 55 mmHg aCSF (Cx43 only), Ca^2+^-free aCSF (Cx43 only) or 50 mM KCl aCSF (Cx46^WT^ and Cx46^QM^ only) each containing 50 μM fluorescein isothiocyanate (FITC) for 10 mins. The cells were then washed, first by perfusion with 20 mmHg in the absence of FITC, before being placed in serial washes of 20 mmHg aCSF. Cells were fixed in 4% paraformaldehyde for 30 mins before being washed 3 times with PBS. Coverslips were mounted inverted on a microscope slide using Fluorshield™ with DAPI mounting medium (Sigma-Aldrich, Cat# F6057). Images were taken using the Zeiss-880 or Zeiss-980 confocal LSM, specifically using the 488 and 561 nm lasers. Subsequent analysis was done using the FIJI software. For Figure 9, figure supplement 1, the median pixel intensity for each transfection was considered to be a statistical replicate.

### Statistical analysis

All quantitative data is presented as box and whisker plots, where the line represents the median, the box is the interquartile range, and the whiskers are the range, with all individual data points included. For multiple comparisons a Kruskal Wallis Anova was performed, followed by post hoc testing via pairwise Mann Whitney *U-*tests (two-tailed) with corrections for multiple comparisons using the false discovery method, with a maximum false discovery set to 0.05 (Curran-Everett, 2000). In text, all data is presented as median (95% CI, upper, lower limit). All calculations were performed on GraphPad Prism.

## Author Contributions

AL: Conceptualisation, data curation, investigation, writing – original draft, review and editing, formal analysis. JB: Investigation (dye loading and confocal microscopy, sensor optimisation). ND: Conceptualisation, supervision, writing – review and editing.

## Data availability

All data is available as supplementary information within this paper.

## Acknowledgements

We thank Prof Alexander Cameron for reading a draft of this paper.

