## Supplemental Figures for "Mechanisms of permselectivity of connexin hemichannels to small molecules"

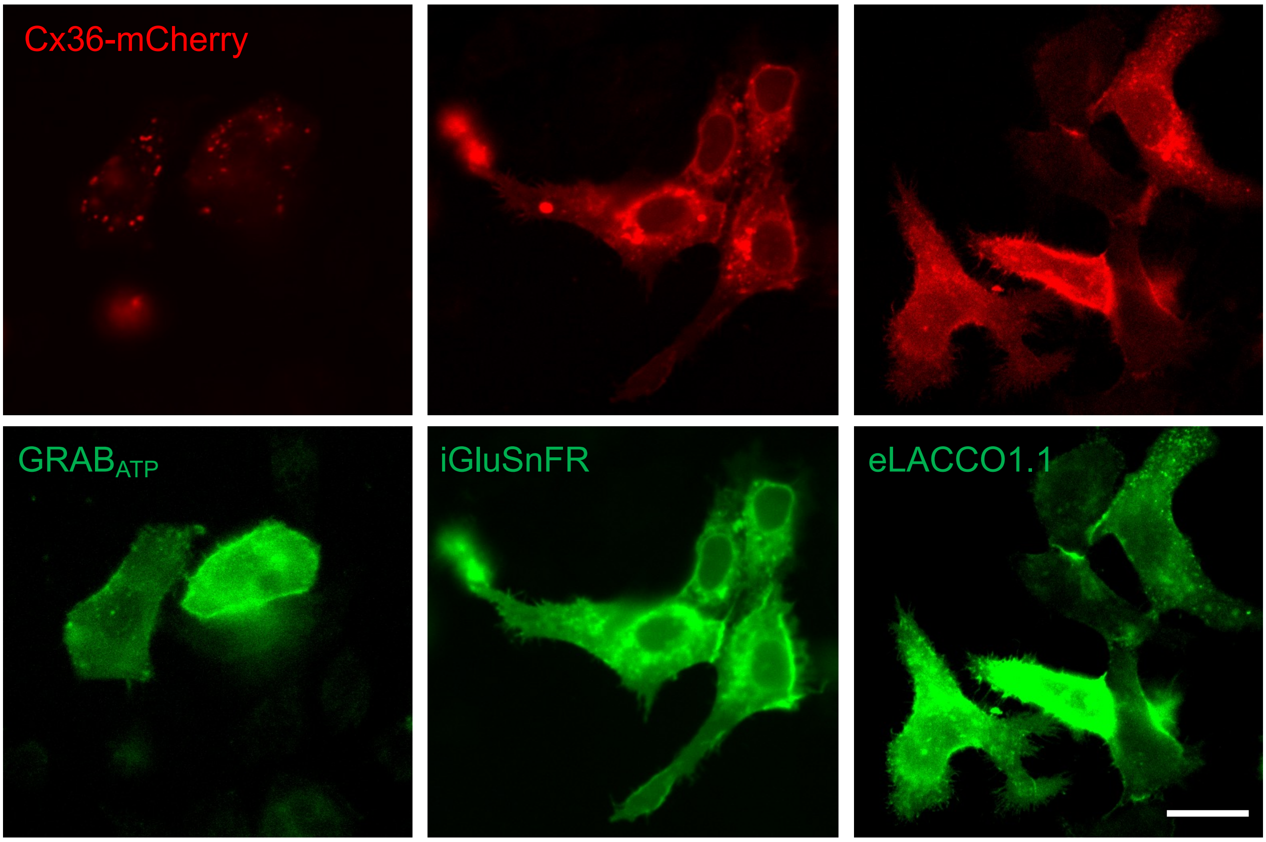


**Supplementary Figure 1. Co-expression of Cx-mCherry and sensor (GFP).** Using Cx36 as an example, the images demonstrate that cells were selected only if they co-expressed both the connexin construct, which is tagged with mCherry (top row), and the sensor, which is tagged with GFP (bottom row). Scale bar represents 20 µm.

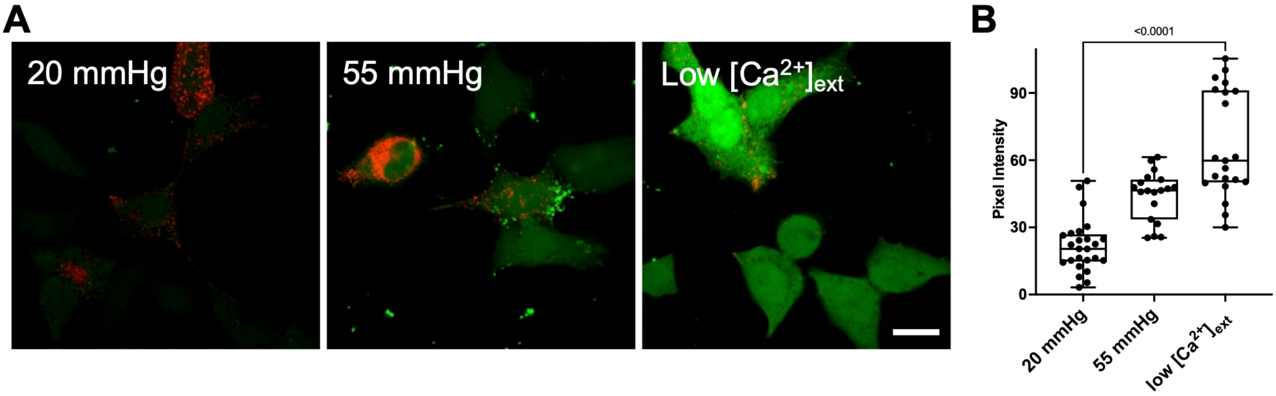


**Supplementary Figure 2. Low $[Ca^{2+}]_{ext}$ is sufficient to open Cx43 hemichannels. A.** Representative images of FITC loaded into mCherry-tagged Cx43 positive cells. Scale bar 20 µm. **B.** Summary data showing the median pixel intensity under control (20 mmHg) (n = 25), hypercapnia (55 mmHg) (n = 19) and low $[Ca^{2+}]_{ext}$ (n = 21). Mann Whitney *U*-test, $p < 0.0001$ (low $[Ca^{2}]_{ext}$ vs 20 mmHg).

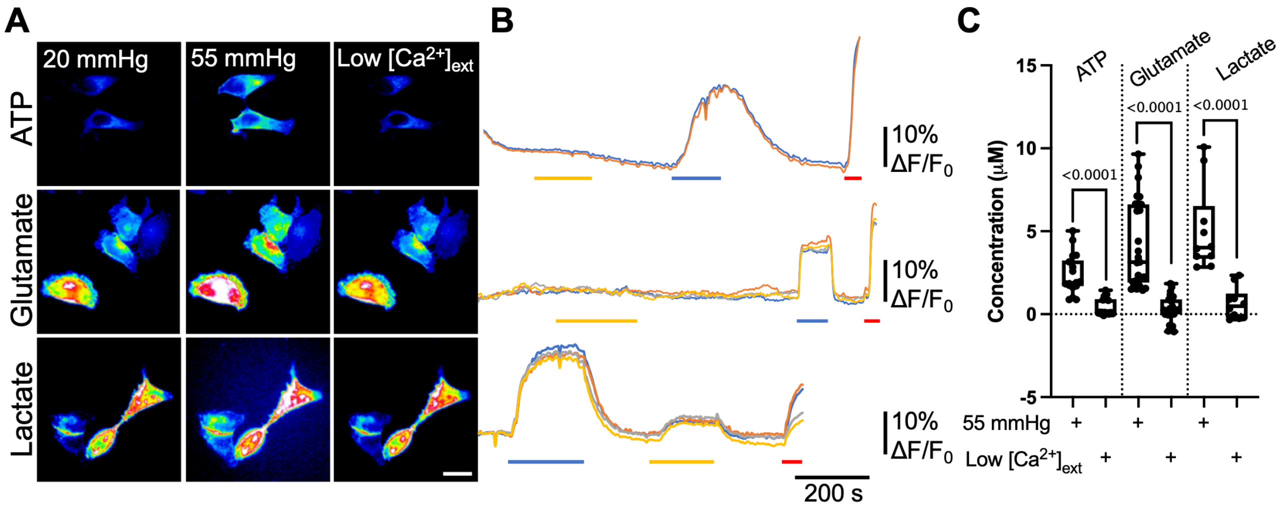


**Supplementary Figure 3. Removal of extracellular $Ca^{2+}$ changes the permeability of Cx43 to physiological molecules. A.** Representative images showing cells from each sensor under each condition. Scale bar represents 20 μm. **B.** Representative traces of normalised fluorescence changes in response to 55 mmHg $pCO_2$ (blue bar), low $[Ca^{2+}]_{ext}$ (yellow bar) and a 3 μM calibration of the corresponding analyte (red bar). **C.** Summary data showing the median release of analytes through Cx43 and how it differs depending on the stimulus: ATP (n = 33 for $CO_2$, n = 17 for high KCl), glutamate (n = 19), lactate (n = 18). Data presented is from at least 3 independent transfections. For 55 mmHg vs low $[Ca^{2+}]_{ext}$ for ATP, glutamate and lactate release, Mann Whitney U-test was performed, $p < 0.0001$.

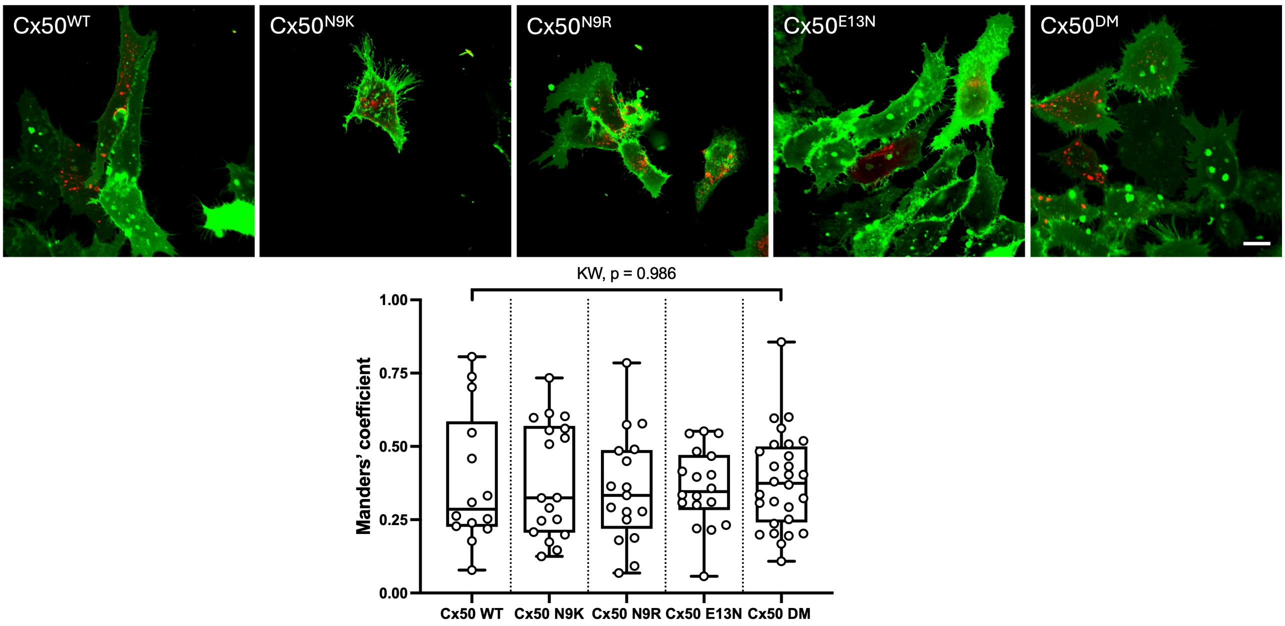


**Supplementary Figure 4. The plasma membrane distribution of Cx50 is unaltered by mutations that alter its permeability to ATP. Top,** Confocal images of DiO staining (green) and the mCherry-tagged Cx50expression (single optical plane). Scale bar 15 μm. $Cx50^{DM}$ is $Cx50^{N9R\ E13N}$. **Bottom,** Manders' analysis of colocalization shows that the proportion of mCherry colocalised with DiO is the same for $Cx50^{WT}$, and the four mutated mutated variants. Data from 3 independent transfections. Kruskal Wallis Anova, p=0.986
